# Longitudinal assessment of mycotoxin co-exposures in exclusively breastfed infants

**DOI:** 10.1101/2020.03.27.011072

**Authors:** Dominik Braun, Eva Schernhammer, Doris Marko, Benedikt Warth

## Abstract

Early-life development of infants may be critically affected by man-made or natural contaminants including mycotoxins. However, data on the occurrence of food contaminants in breast milk is scarce and prohibits a comprehensive exposure and risk assessment for mothers and their infants.

Here, we present a longitudinal exposure assessment over the first 211 days of a single newborn girl (study A) by measuring multiple mycotoxins in milk. Eighty-seven consecutive breast milk samples were obtained from the newborn’s mother living in Austria and following a regular mixed diet. Mycotoxins were analyzed by utilizing a highly sensitive LC-MS/MS approach covering 29 mycotoxins and key metabolites. In addition to this longitudinal study, three mothers provided breast milk samples each on five consecutive days, for a preliminary comparison of inter-day and inter-individual variation in exposures (study B). Study A revealed that mycotoxin occurrence in breast milk was limited to the emerging mycotoxins alternariol monomethyl ether (AME), beauvericin (BEA), enniatins (A, A_1_, B, B_1_) and to ochratoxin A (OTA), which is regulated in commercial infant food. These mycotoxins were, if present, mostly detected at very low concentrations (<10 ng/L), except AME which exceeded this concentration on two distinct days by a factor of 3x and 5x. Overall, longitudinal results indicated chronic low-dose exposure to the detected mycotoxins. Other regulated mycotoxins including the carcinogenic aflatoxins or the estrogenic zearalenone and their biotransformation products were absent in all tested samples. Study B confirmed the results of study A, with minimal inter-day and inter-individual variation. In addition, a preliminary correlation of OTA levels occurring in breast milk and matched urine samples was found (r=0.64, p=0.034) in study B. Based on the data set obtained in study A, exposure of the infant was estimated. Exposure estimates of individual mycotoxins were on average below 1 ng/kg body weight per day.

Our preliminary findings suggest that recommended maximum daily intake levels might not be exceeded in the Austrian population. However, exposure is likely to be higher in populations with lower food safety standards. In the light of co-occurrence of several emerging mycotoxins in breast milk, future studies should address low-dose mixture effects. This also includes other environmental contaminants which may be present in this bio-fluid and should involve an exposome-scale risk assessment. All these efforts must be intended to minimize exposure of mothers and infants in a window of high susceptibility.

## Introduction

Mycotoxins are naturally occurring toxic secondary metabolites produced by various fungi, including *Aspergillus, Fusarium, Penicillium* and *Alternaria* species. Toxigenic fungi frequently contaminate agricultural crops pre- or postharvest in diverse environmental conditions (Bennett and Klich 2003). The ubiquitous occurrence of mycotoxins in the food chain has been shown in numerous reports over the last decades (Eskola *et al*. 2019; Schatzmayr and Streit 2013). Several *in vivo* studies documented harmful effects including immune suppression, target organ toxicity, genotoxicity or carcinogenicity. As a result, legislative regulations were introduced for the major mycotoxins and maximum tolerated limits (MTL) were implemented to control, measure and diminish their occurrence (EC 2006). Mycotoxins of legislative interest are aflatoxin B_1_ (AFB_1_), AFB_2_, AFG_1_, AFG_2_ and AFM_1_ due to their known carcinogenic effect. In addition to cancer, aflatoxins have also been linked to growth stunting in children and the suppression of immune responses (Gong *et al*. 2004; Gong *et al*. 2012; *Turner *et al*. 2003). Of Fusarium* derived mycotoxins, trichothecene deoxynivalenol (DON) has been shown to cause emesis and to inhibit protein biosynthesis (Beasley 2017; EFSA and Knudsen 2017). Zearalenone (ZEN) is known for its ability to bind to the estrogen receptor (Kowalska *et al*. 2018). Ochratoxin A (OTA) critically affects the kidney due to accumulation in the nephron (Malir *et al*. 2013), while exposure to fumonisins has been linked to esophageal cancer (Shirima *et al*. 2015). Despite their documented adverse effects, to date, few mycotoxins are regulated and monitored. In addition, so-called emerging mycotoxins like *Fusarium*-derived enniatins and beauvericin (BEA) or mycotoxins produced by *Alternaria*, e.g. alternariol and its monomethyl ether (AOH and AME) are frequently detected in various food matrices (Puntscher *et al*. 2019). Most of these fungal secondary metabolites are toxicologically not fully characterized or data on their occurrence is lacking, hence, hindering proper risk assessment. The concept of ‘developmental origins of health and disease’ (DOHaD) postulates that early-life exposures affect later-life health outcomes (Barker 2000; Bateson *et al*. 2004; Mandy and Nyirenda 2018). In general, infants are more vulnerable to the risks of mycotoxins as their detoxification capabilities are not fully developed and their intake of food is comparatively higher. In this critical time window of infancy, the World Health Organization (WHO) recommends to exclusively breastfeed newborns and infants for at least six months (Horta 2007). Breastfeeding is associated with numerous positive effects on both mother and child such as emotional bonding, tailored nutritional diet, uterine involution and the reduction of breast cancer risk (Jäger *et al*. 2014; Palmer *et al*. 2014). However, it may also pose unexpected risks: maternal exposure to food contaminants, such as mycotoxins, for example, may subject offspring not only to *in utero*, but also breast milk mycotoxins, thereby potentially increasing the risk for childhood and adulthood disease through gene x environment interactions or adverse epigenetic programing (Thornburg *et al*. 2010; Warth *et al*. 2019). Currently, no clear data on excretion patterns of food contaminants in breast milk at different stages during breastfeeding exist (LaKind *et al*. 2018; Lehmann *et al*. 2018). To date, studies have mainly focused on AFM_1_ and OTA, and they used enzyme linked immunosorbent assays (ELISA) or liquid chromatography coupled to fluorescence detection (LC-FD), both of which lack the specificity and sensitivity of state-of-the-art LC-MS systems as reviewed by Warth *et al*. (2016) and Sengling Cebin Coppa *et al*. (2019). Recently, we developed and validated two targeted LC-MS/MS assays to simultaneously assess multiple classes of mycotoxins in breast milk. The methods proved to be fit for purpose and allow the evaluation of these natural contaminants down to the pg/L range, enabling quantitative exposure assessment for infants even in countries with high food safety standards and low mycotoxin risk (Braun *et al*. 2018; Braun *et al*. 2020).

Based on this technological progress, the primary aims of the presented experiments were a) to obtain preliminary longitudinal occurrence data of multiple mycotoxins in breast milk (study A) for estimating infant exposure, which is subsequently compared to established guidance values and b) to evaluate inter-day and inter-individual differences of mycotoxin co-occurrence patterns in a proof-of-principle experiment (study B).

## Materials and methods

### Chemicals and Reagents

Acetonitrile (ACN), methanol (MeOH) and water were purchased from Honeywell (Seelze, Germany). For sample preparation, ammonium acetate, formic acid, anhydrous magnesium sulfate, sodium chloride and formic acid were purchased from Sigma Aldrich (Vienna, Austria). The following toxins were purchased as reference material: Aflatoxin B_1_ (AFB_1_), AFB_2_, AFG_1_, AFG_2_, deoxynivalenol (DON), OTA, nivalenol (NIV), Sterigmatocystin (STC), fumonisin B_1_ (FB_1_), FB_2_, T-2 toxin, alpha zearalenol (α-ZEL), β-ZEL, alpha zearalanol (α-ZAL), β-ZAL, zearalanone (ZAN) and ZEN from RomerLabs (Tulln, Austria); and aflatoxin metabolites AFM_1_, AFM_2_, AFP_1_, AFQ_1_, AFB_1_-N7-guanine adduct, as well as AME, AOH, beauvericin (BEA), Citrinin (CIT), HT-2 toxin, ochratoxin alpha (OTα), ochratoxin B (OTB) from Toronto Research Chemicals (Ontario, Canada). Enniatin A (Enn A), Enn A_1_, Enn B, Enn B_1_ and TEN were purchased from Sigma-Aldrich (Vienna, Austria). Dihydrocitrinone (DH-CIT) was kindly provided by Prof. Michael Sulyok (IFA-Tulln, Austria). Solid reference standards were dissolved in ACN to final concentrations of 5-500 μg/mL as stock solutions. AFB_1_-N7-guanine was dissolved in ACN/H_2_O/acetic acid (75/24/1, v/v/v) according to manufacturer’s instructions. The multi-toxin working solution containing all analytes was prepared by diluting individual stock solutions to concentrations of 36 - 17,000 ng/mL. ^13^C-labelled reference standards of AFM_1_, CIT, DON, NIV, OTA and ZEN were purchased from RomerLabs (Tulln, Austria). A fresh mix of internal standards was prepared regularly reaching final concentrations of 0.1 - 4.5 ng/mL. All solutions and solid reference standards were stored at −20 °C.

### Sample collection

Human milk samples from two different experiments were investigated. In study A, longitudinal samples (n = 87) were provided by a lactating mother between July 2015 and January 2016. Breast milk was pumped into sample containers at the mother’s home. Multiple samples collected at the same day, partly over two days, were combined to an aggregated sample. This was performed depending on the needs of the mother and her infant, and left-overs not consumed by the infant were mixed after at least two days during which the samples were stored at 4 °C. Thus, the mixed sample might not have been blended with the same volume of each meal. Although possible variability in occurrence of mycotoxin concentration remains due to the dynamic nature of this bio-fluid, this approach was deemed most suitable to obtain representative samples. After combining multiple meal left-overs, samples were kept at −20 °C until analysis. On some days all breast milk was consumed by the infant leaving no sample for laboratory analysis. In study B, three mothers collected samples in the morning by manually expressing milk into containers at home each day on five consecutive days. In addition, first morning urine samples were collected over the five-day period. All collected breast milk and urine samples were immediately stored at −20 °C. Sampling was conducted about one to three months after delivery. Socio-demographic data for all mothers were recorded. Their age ranged between 25-32 years with a normal body mass index. Two participants were primiparous women, two had a high level of education (university degree), while all mothers earned a medium income. Throughout both studies all mothers maintained their regular mixed diet, thus, none was following vegetarian, vegan or special diet plans. The study was approved by the Ethics Committee of the government of Lower Austria and the University of Vienna Ethics Committee under the authorization number #00157.

### Sample preparation protocol and LC-MS/MS analysis

For breast milk mycotoxin analysis, the optimized protocol of Braun *et al*. (2020) was used. In brief, samples were thawed, homogenized and 1 mL breast milk sample was extracted with 1 mL of acidified ACN (1% formic acid). After homogenization, anhydrous magnesium sulfate (0.4 g) and sodium chloride (0.1 g) were separately added and mixed. The upper layer (ACN, 950 μL) was transferred into a micro-reaction tube after a centrifugation step (4,750 x g, 10 min, 4 °C). This extract was chilled at −20 °C for 2 h following a second centrifugation step (14,000 x g, 2 min, 4 °C). The supernatant (900 μL ACN extract) was directly diluted in a H_2_O preloaded reservoir to 5% ACN and subsequently loaded to an Oasis PRiME HLB^®^ solid phase extraction (SPE) column (1cc, 30 mg, Waters, Milford, MA). This column was equilibrated with 1 mL ACN and conditioned with 1 mL H_2_O/ACN (95/5, v/v) prior to sample loading. Following sample loading, the column was washed twice (5%ACN, 500 μL) and mycotoxins were eluted using neat ACN (three times 500 μL). This extract was evaporated to dryness using a vacuum concentrator (Labconco, Missouri, USA) and samples were reconstituted in 81 μL MeOH/ACN (50:50, v:v) and 9 μL of the internal standard mixture. After homogenization and ultra-sonication (5 min), the samples were transferred to amber LC-vials and analyzed.

Details of the LC-MS/MS method to quantify mycotoxins in breast milk, including instrumental parameters and analytical figures of merit were reported in Braun *et al*. (2020), while a brief summary of the methods’ basic performance parameters are reported in Table S1. In brief, the LC-MS/MS system consisted of an Agilent 1290 Infinity II LC system coupled to a Sciex QTrap^®^ 6500^+^ mass spectrometer (Darmstadt, Germany). The MS was equipped with a Turbo-V™ electrospray ionization interface. Optimized chromatographic performance was achieved using an Acquity HSS T3 column (1.8 μm, 2.1×100 mm) guarded with a VanGuard pre-column (1.8 μm, Waters, Vienna, Austria) at a flow rate of 0.25 mL/min. A binary gradient elution was used consisting of an acidified ammonium acetate solution in water (5mM, acidified with 0.1% acetic acid; A) and MeOH (B). To baseline-separate all analytes of interest, a multi-step gradient was applied as follows: Methanol was kept at 10% for the first 0.5 min, before increasing it to 35% within half a minute. Then, eluent B was linearly raised to 60% until 3.0 min and to 97% until 10.0 min. After 6.0 min at 97% eluent B, the column was re-equilibrated using initial conditions (10% B) between 16.1 and 19.0 min, resulting in a total runtime of 19 min. The instruments’ autosampler and column oven were maintained at 10 °C and 40 °C, respectively. For analysis, a sample volume of 3 μL was injected onto the column. The MS was acquiring in scheduled multiple reaction monitoring (sMRM) applying fast polarity switching. The sMRM algorithm was used to limit the measurement of analytes around their expected retention time and thus increase dwell time and reduce the overall cycle time.

For urinary analysis, mycotoxins were extracted following the protocol of Šarkanj *et al*. (2018). A brief summary of the extraction procedure is included in the Supplementary Material. Details of the LC-MS/MS method to quantify mycotoxin in urine, including instrumental parameters (Table S2) and performance parameters during in-house validation (Table S3) are reported in the Supplementary Material as well. Analyst^®^ (version 1.7) software was used for data acquisition and instrument control. Data was processed using the Sciex OS software package (version 1.6).

### Daily intake estimation

The estimated daily intake (EDI) of mycotoxins via breast milk as reported in Table S4, was based on an upper bound deterministic approach using the longitudinal exposure data from study A. Mycotoxin exposures via milk formula or cereal based products could be excluded, as the infant was exclusively breastfed. All mycotoxins detected below their respective LOQ values were set to half the LOQ value (LOQ/2), while the concentration of not detected mycotoxins were assigned to half the LOD value (LOD/2), respectively. The studied time frame of 211 days was subdivided into seven different intervals. The first weeks after delivery were sectioned into four intervals of 10 days to account for a linear increase in milk intake (Table S4). Segment five included the first quarter following the second quarter in segment six. Interval 7 consisted of the last sample time point on day 211 after delivery. The infant’s body weight (bw) was recorded by a pediatrician during regular, prophylactic mother- and-child examinations. Breast milk intake was estimated on several occasions by pumping milk into containers with volume levels and recording the consumed volume after the infant was fed. Infant’s bw as well as the daily breast milk intake volume (DBI) were assumed to be constant in the 10 day segments, while bw and DBI was averaged in the remaining time intervals. The EDI (ng/kg bw) was calculated by multiplying the concentration of the respective mycotoxin and the DBI, which was then divided by the body weight (EC 2002).

### Statistical analysis

Data analysis was performed using MS Excel 2019 and OriginPro 2020 (v9.7). Descriptive statistics were derived for all samples with detectable mycotoxin concentration (>LOD). Spearman correlation was performed for mycotoxins with a detection rate of >80% (i.e above the LOD) and was calculated between co-occurring mycotoxins in study A and B, respectively. In addition, Spearman correlation was calculated for mycotoxins which simultaneously occurred in breast milk and urinary samples in study B.

## Results and discussion

### Longitudinal mycotoxin exposure during early-life

Breast milk samples obtained of a single volunteer during a timeframe of 211 days after delivery (n=87) were tested for 29 mycotoxins and their key metabolites applying a highly sensitive and selective LC-MS/MS multianalyte method in study A. To the best of our knowledge, this is the first study to examine longitudinal concentration changes of multiple mycotoxins in breast milk. Occurrence data of environmental contaminants in this bio-fluid are insufficiently understood and little is known about the transfer of most mycotoxins into breast milk in the recommended breastfeeding timeframe (Lehmann *et al*. 2018). However, potential mycotoxin levels were expected to be very low, as high food safety standards are enforced in the European Union. Our results obtained in this preliminary experiment shed a detailed picture on co-exposures and excretion patterns. As expected, mycotoxins that were identified were quantified only in the low ng/L-range. Detailed descriptive statistics on quantified mycotoxins are reported in Table 1. Mycotoxins detected during the investigated period included the rather lipophilic AME, BEA, EnnA, EnnA_1_, EnnB, EnnB_1_ and OTA (Figure 1).

**Table 1.**
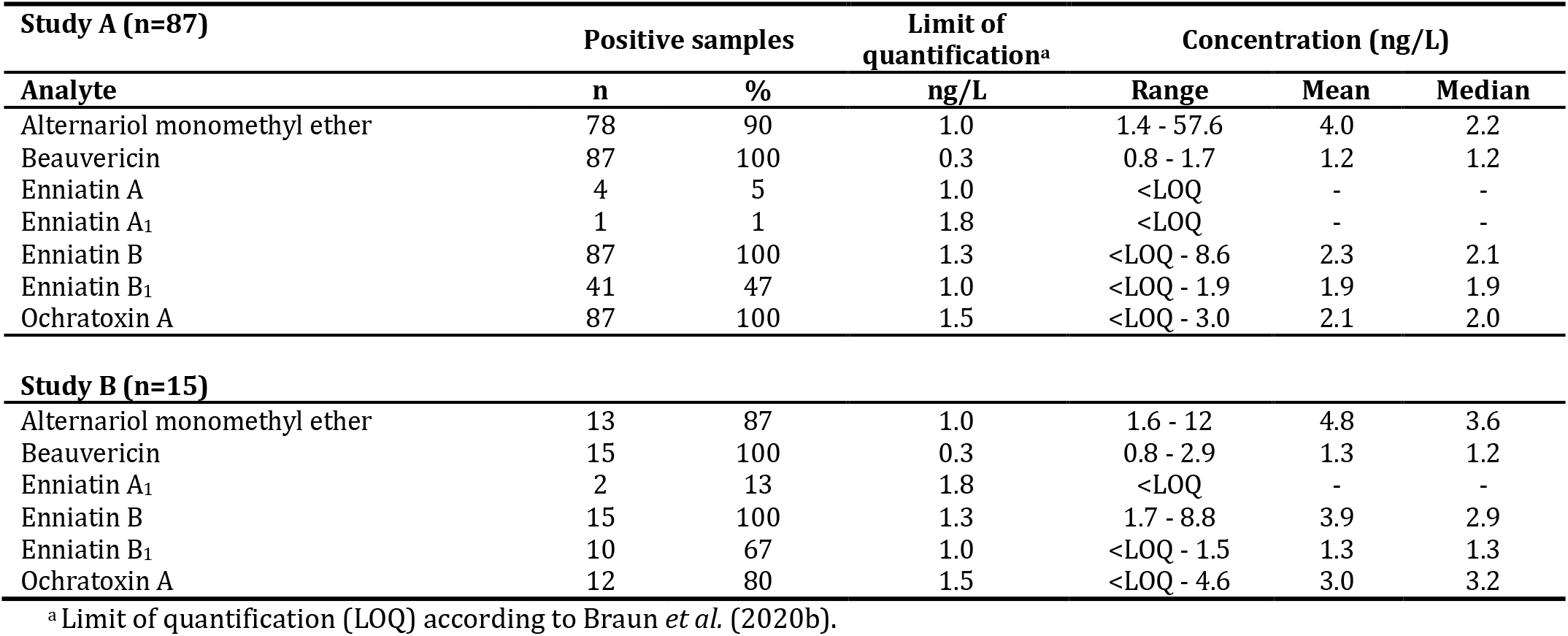
Detailed descriptive statistics of mycotoxins detected in breast milk samples (n=87) obtained from a volunteer within 211 days after delivery (study A) and in breast milk samples (n=15) obtained from three volunteers on five consecutive days (study B).

| Study A (n=87) | Positive samples |  | Limit of quantification<sup>a</sup> | Concentration (ng/L) |  |  |
| --- | --- | --- | --- | --- | --- | --- |
| Analyte | n | % | ng/L | Range | Mean | Median |
| Alternariol monomethyl ether | 78 | 90 | 1.0 | 1.4 - 57.6 | 4.0 | 2.2 |
| Beauvericin | 87 | 100 | 0.3 | 0.8 - 1.7 | 1.2 | 1.2 |
| Enniatin A | 4 | 5 | 1.0 | <LOQ | - | - |
| Enniatin A<sub>1</sub> | 1 | 1 | 1.8 | <LOQ | - | - |
| Enniatin B | 87 | 100 | 1.3 | <LOQ - 8.6 | 2.3 | 2.1 |
| Enniatin B<sub>1</sub> | 41 | 47 | 1.0 | <LOQ - 1.9 | 1.9 | 1.9 |
| Ochratoxin A | 87 | 100 | 1.5 | <LOQ - 3.0 | 2.1 | 2.0 |

| Study B (n=15) | Positive samples |  | Limit of quantification<sup>a</sup> | Concentration (ng/L) |  |  |
| --- | --- | --- | --- | --- | --- | --- |
| Alternariol monomethyl ether | 13 | 87 | 1.0 | 1.6 - 12 | 4.8 | 3.6 |
| Beauvericin | 15 | 100 | 0.3 | 0.8 - 2.9 | 1.3 | 1.2 |
| Enniatin A<sub>1</sub> | 2 | 13 | 1.8 | <LOQ | - | - |
| Enniatin B | 15 | 100 | 1.3 | 1.7 - 8.8 | 3.9 | 2.9 |
| Enniatin B<sub>1</sub> | 10 | 67 | 1.0 | <LOQ - 1.5 | 1.3 | 1.3 |
| Ochratoxin A | 12 | 80 | 1.5 | <LOQ - 4.6 | 3.0 | 3.2 |

<sup>a</sup> Limit of quantification (LOQ) according to Braun *et al.* (2020b).

**Figure 1.**
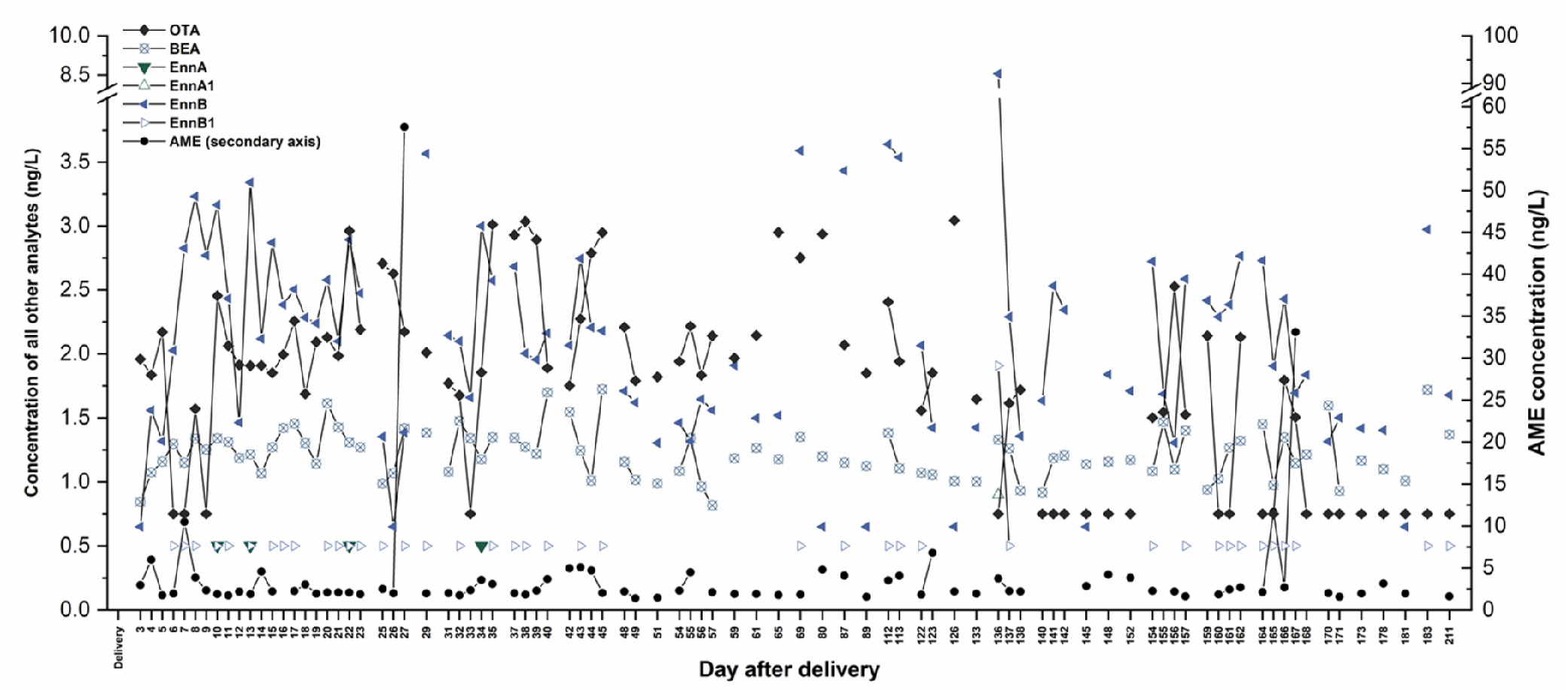
Dynamic exposure of mycotoxins in breast milk (study A): alternariol monomethyl ether (AME, ●), beauvericin (BEA, ⊗), enniatin A_1_ (EnnA, ▼), enniatin A_1_ (EnnA_1_, △), enniatin B (EnnB, ◀), enniatin B_1_ (EnnB_1_, ▷) and ochratoxin A (OTA, ◆) occurring in breast milk between days 3 and 211 postpartum (n=87). Line connected points indicate the changes on consecutive days. No linear time scale was chosen to plot mycotoxin concentration data of all samples in one graph. Detected analytes which were below the LOQ value were set to half the LOQ (LOQ/2). Other analytes, e.g. aflatoxin M_1_ (AFM_1_; LOQ: 4.0 ng/L), zearalenone (ZEN; LOQ: 32 ng/L) and citrinin (CIT; LOQ: 6.0 ng/L) were not detected in any sample.

Quantification of AME was possible in 90% of the analyzed samples with a maximum concentration of 58 ng/L on day 27 postpartum. AME was quantified below 10 ng/L in most evaluated samples with the exception of day 7 (11 ng/L), 165 (12 ng/L) and 167 (33 ng/L) postpartum, respectively. Quantification of BEA was possible in all samples (100%) with a highly constant concentration range between 0.8 to 1.7 ng/L. The same was true for EnnB with a mean concentration of 1.9 ng/L, although on some days the concentration was below the respective LOQ value. EnnB_1_, which differs to EnnB only by the presence of an isobutyl residue instead of an isopropyl residue, was found in 47% of the analyzed samples. However, except on day 136 postpartum (1.9 ng/L) EnnB_1_ concentrations were below the LOQ value (Figure 2). On days 136, 137 and 138 postpartum the concentration of EnnB and EnnB_1_ decreased simultaneously. Contrarily, on these days the OTA levels increased up to 1.7 ng/L. In 36 out of 87 analyzed samples up to five mycotoxins were detected, while in most samples (90%) at least four mycotoxins, namely AME, BEA, EnnB and OTA co-occurred. Spearman correlation analysis for these four toxins revealed that BEA and EnnB were modestly, although highly significant associated (r=0.43; p=7.4E-6; Figure 3A), which might be partly explained due to their structural similarity. Other mycotoxin combinations were not associated in any way.

**Figure 2.**
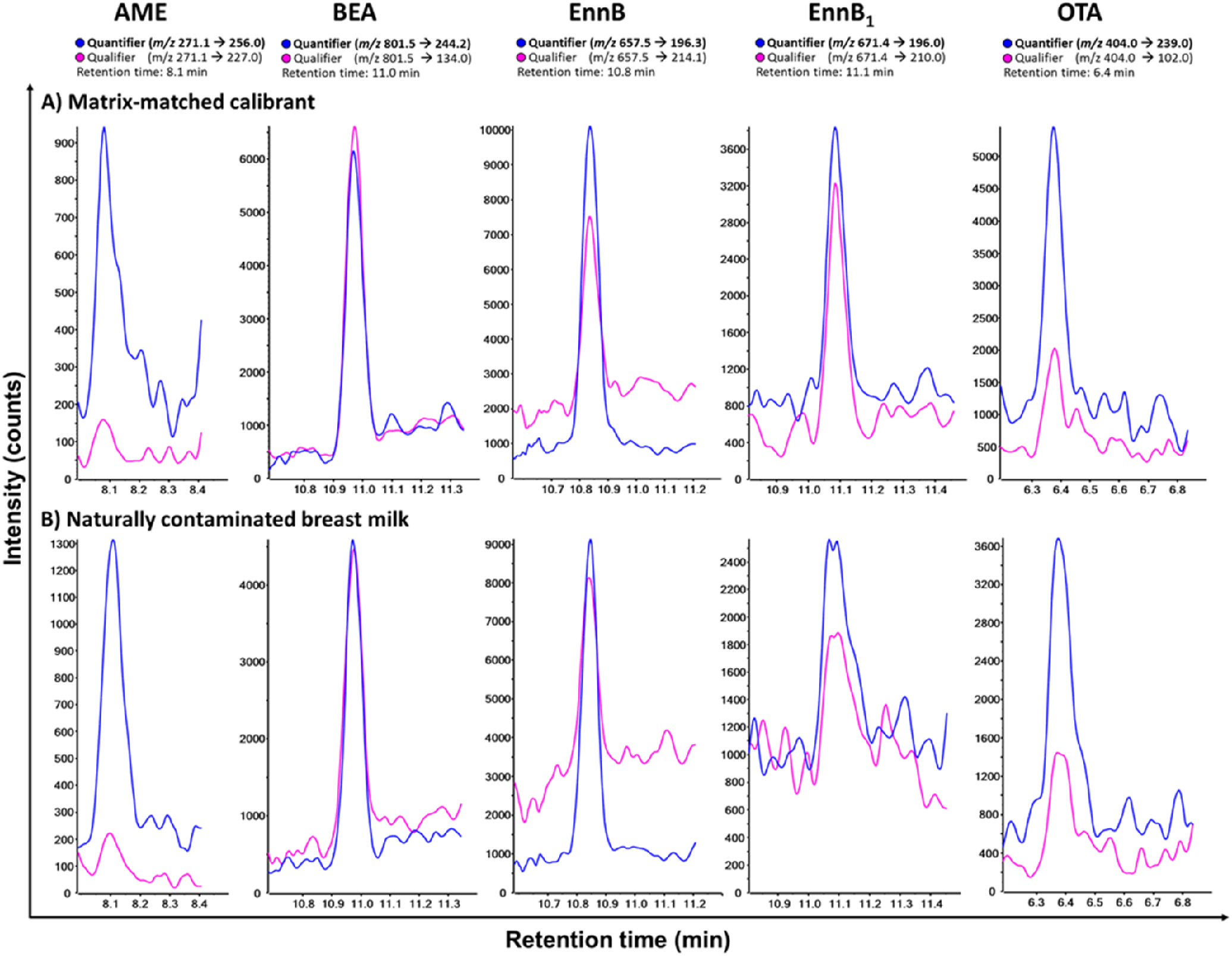
LC-MS/MS chromatograms demonstrated mycotoxin co-exposure in the Austrian breast milk sample on day 35 after delivery: alternariol monomethyl ether (AME; 3.1 ng/L), beauvericin (BEA; 1.3 ng/L), enniatin B (EnnB; 2.6 ng/L), enniatin B_1_ (EnnB_1_: <LOQ) and ochratoxin A (OTA; 3.0 ng/L). For all toxins a fortified matrix-matched calibrant (A) and the Austrian breast milk sample (B) are shown, respectively

**Figure 3.**
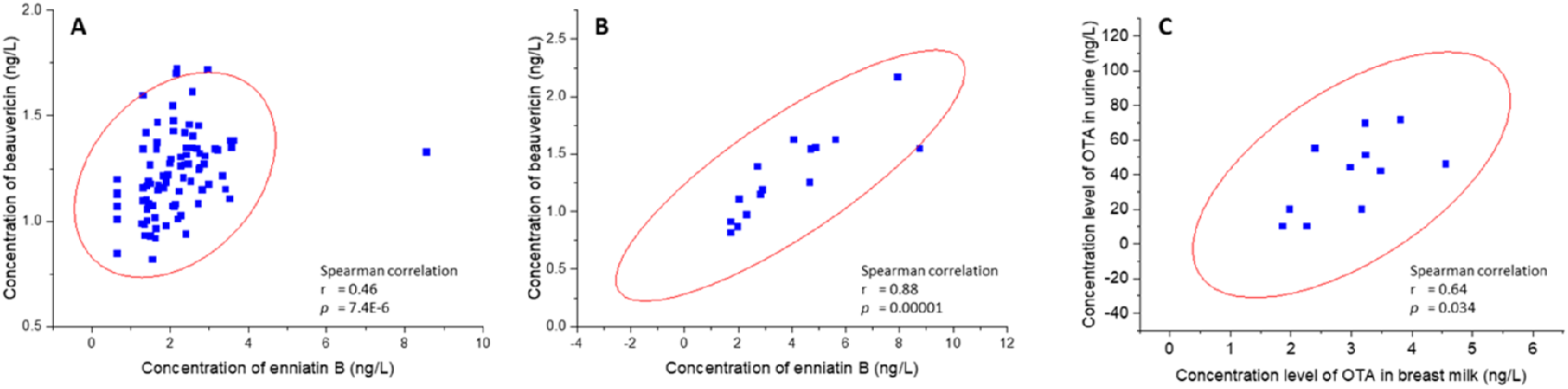
Relationship of beauvericin and enniatin B concentration levels in breast milk in study A (A) and study B (B) respectively. In addition, relationship of ochratoxin A (OTA) concentration levels in breast milk and urine as obtained in study B (C).

With regards to the concentration range, our results are comparable to others who studied the occurrence of multiple mycotoxins. Andrade *et al*. (2013) analyzed aflatoxins and OTA in 224 samples obtained from different human milk banks in Brazil using LC-FD. The authors stated that OTA was not detectable in any sample with an LOQ value of 10 ng/L (Andrade *et al*. 2013). Tonon *et al*. (2018) reported a LOQ value of 12.5 ng/L using LC-MS/MS and did not detect OTA in any sample. Even though in the present study OTA was detected in all samples and quantification was possible in 74% of analyzed samples, the maximum concentration of OTA was 3 ng/L, again, highlighting the ultimate sensitivity of the method applied in the study at hand.

Others measured breast milk OTA levels in Chilean mothers on five different occasions over the time course of four months. In contrast to our preliminary experiments described here, the authors reported a mean concentration of 52 ng/L and the highest measured concentration was observed during the first six days after delivery (Munoz *et al*. 2014). Clearly, influencing factors such as geographical region, seasonal changes or number of births may have an enormous impact on the composition of breast milk which affects mycotoxin occurrence patterns. In addition, most of the measured toxins are rather lipophilic and may be remobilized out of adipose tissue during lactation (LaKind *et al*. 2009) or, as it is described for OTA, have a prolonged binding to plasma albumin (Il’ichev *et al*. 2002; Studer-Rohr *et al*. 2000). Thus, even if a mother was not recently exposed to OTA via her diet, the prolonged half-life through protein binding affect its availability for lactational transfer.

Recently, similar mycotoxin concentration levels as obtained in study A were observed in 22 breast milk samples obtained from Nigerian mothers. Those results were measured using the same methodology applied here, however, only a single time point was evaluated. In the former study, AME, BEA and OTA were the most frequently found toxins with concentrations up to 11, 12 and 68 ng/L, respectively (Braun *et al*. unpublished, Ezekiel *et al*. unpublished). EnnB and EnnB_1_ were detectable at lower concentrations (Braun *et al*. 2018; Braun *et al*. 2020).

In contrast to these results, high mycotoxin levels were reported by Rubert *et al*. (2014) in breast milk obtained from Spanish mothers. Using LC coupled to a high-resolution mass spectrometer the authors reported e.g. EnnB and EnnB_1_ in two out of 21 samples at a mean concentration of 105,000 ng/L and 96,000 ng/L, respectively. In addition, ZEN was reported in 62% of the samples in a concentration range of 2,100 to 14,000 ng/L (Rubert *et al*. 2014), whereas Austrian or Nigerian samples were not contaminated with ZEN (LOD 16 ng/L) (Braun *et al*. 2020). Hence, it is not unlikely that the Spanish study over-estimated exposure levels as no internal standards were applied.

Recently, the occurrence of very low levels of AME and BEA in breast milk was reported for the first time in a limited number of Nigerian, but also in Austrian breast milk (Braun *et al*. 2020). In the current pilot study, we could verify our findings and additionally observed chronic low background exposures. Interestingly, at day 7, 27, 165 and 167 postpartum the concentration of AME increased above a concentration of 11 ng/L. and declined during the following two days. The sudden concentration change in breast milk can most likely be traced back to contaminated food products, while the possible release of this rather lipophilic compounds out of adipose tissue might be the reason for the chronic low-level occurrence in the studied time interval. Here, dietary recalls and food analysis may give detailed insights for addressing these differences in follow-up studies. AME is frequently detected in European food items, e.g. in tomato sauce, sunflower seed oil, wheat flour or strawberries in a concentration range of 0.06 - 4.7 ng/g (Juan *et al*. 2016; Puntscher *et al*. 2019). In contrast to AME, BEA did not exhibit any peak occurrence patterns during the investigated time frame. The small concentration range and the presence of this mycotoxin in all samples suggest chronic background exposures through the diet and spot sampling is likely sufficient to obtain representative exposure data for the detected toxins. BEA is a common contaminant found in cereal based products such as wheat, corn or rice in a concentration range of 0.03 - 10 μg/kg (Covarelli *et al*. 2015; Jestoi *et al*. 2004), indicating continuous low-level exposure via food. Thus, it is likely that even with a variable diet chronic background exposure may not be avoided. Overall, the analyte concentration of the detected toxins was rather stable over the studied time period. Other mycotoxins, e.g. aflatoxins, CIT, OTB, OTα, ZEN, ZAN or their key metabolites were not detected in any sample above their respective LOD value (Table S1) in this experiment.

To evaluate the risk for infants, hazard information on mycotoxins reported in this pilot study and the resulting exposure (see section 0) have to be assessed. The emerging *Fusarium* toxins BEA and enniatins mediate low acute toxicity *in vivo*, exhibit ionophoric properties which lead to a dysfunction of mitochondria and lysosomes, and are able to induce aggregation of blood platelets above a concentration of 20 μmol/L (Jestoi 2008; Oliveira *et al*. 2020; Tonshin *et al*. 2010). The Alternaria metabolite AME has received more attention in recent years due to its genotoxic properties (Ostry 2008). At micromolar concentrations (approximately 300 μg/L and higher) it has been shown *in vitro* that AME induces DNA strand breaks by topoisomerase poisoning (Fehr *et al*. 2009; Schwarz *et al*. 2012). In addition, weak estrogenic effects *in vitro* and effects on the reproductive performance of pigs and other mammals were postulated (Dellafiora *et al*. 2018; Tiemann *et al*. 2009). OTA has been investigated over the last decades in numerous studies and reviews (Haighton *et al*. 2012; O’Brien and Dietrich 2005). Key adverse effects include renal toxicity and carcinogenesis (Limonciel and Jennings 2014). In addition, neurotoxic, teratogenic and hepatotoxic effects were described in animal models and *in vitro* (Doi and Uetsuka 2011; Malir *et al*. 2013).

### Inter-individual mycotoxin exposure on five consecutive days in breast milk and urine

To gain further insights into mycotoxin occurrence patterns, breast milk and matched urine samples of three study subjects living in Austria were analyzed on five consecutive days (study B). The results obtained are illustrated in Figure 4. Mycotoxins identified in these 15 breast milk samples included AME, BEA, EnnA_1_, EnnB, EnnB_1_ and OTA. Similar to study A, all other tested mycotoxins were not present in any sample above the LOD value (Table S1). All analyzed samples were contaminated with BEA and EnnB with maximum concentrations of 2.2 and 8.8 ng/L, respectively. The maximum concentration of 12 ng/L was detected for AME, which is in agreement with the longitudinal data. In fact, comparing the results discussed above (study A; Table S5) and the data retrieved within five days (study B; Table S6) the proof-of-principle experiments had a similar outcome. Also, EnnB_1_ and OTA occurrence with 67% and 80% were in the range of the longitudinal detection frequency. Average and median concentration levels of individual mycotoxins were comparable between the three subjects. Five mycotoxins co-occurred in all breast milk samples analyzed from two mothers, whereas at least two and up to four mycotoxins were present simultaneously in the samples obtained from the third mother. Similar to study A, spearman correlation analysis revealed that BEA and EnnB were significantly associated (r=0.88, p= 0.00001; Figure 3B) in this proof-of-principle experiment.

**Figure 4.**
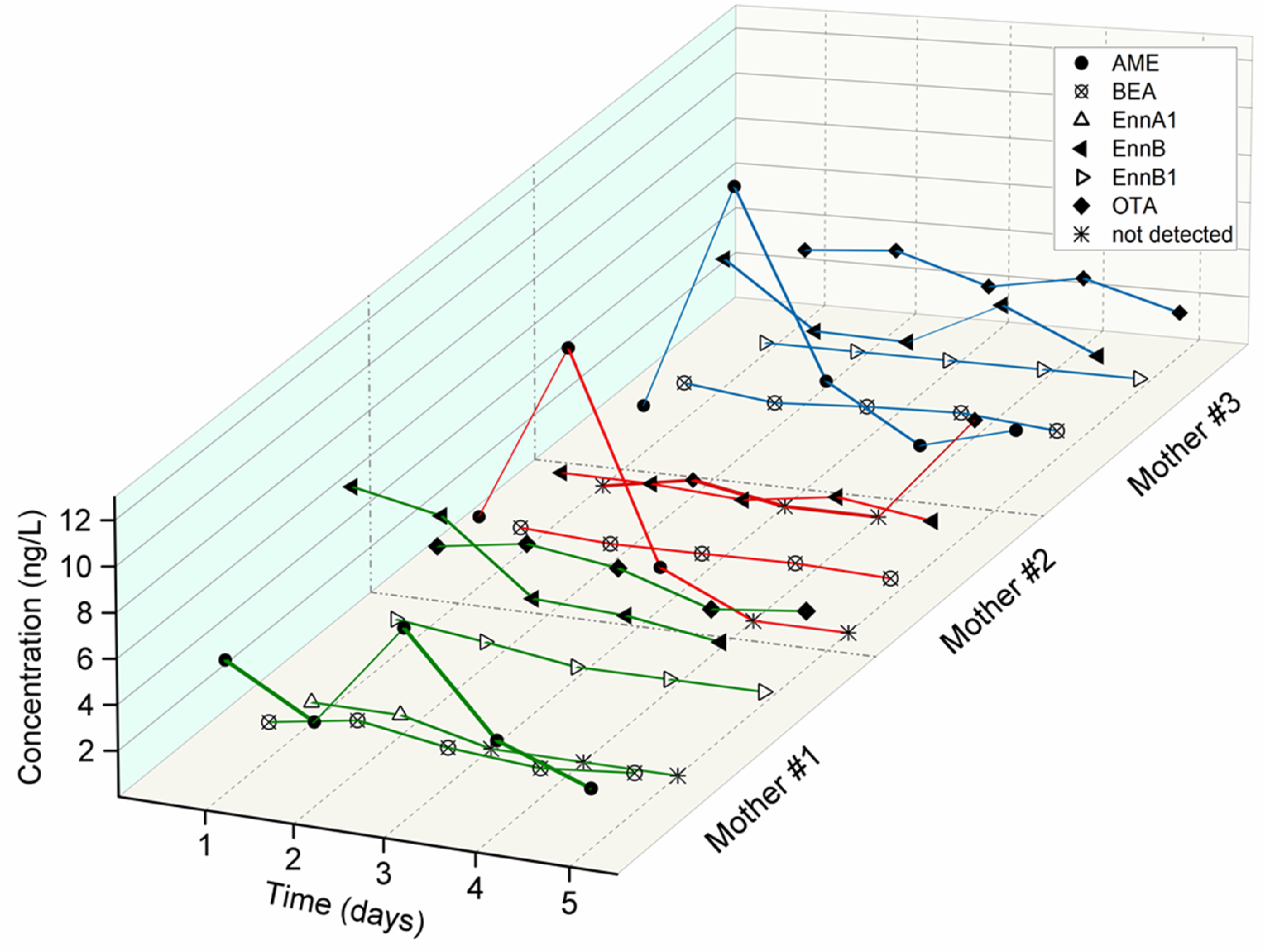
Dynamic exposure of mycotoxins in breast milk (study B): alternariol monomethyl ether (AME, ●), beauvericin (BEA, ⊗), enniatin A_1_ (EnnA_1_, △), enniatin B (EnnB, ◀), enniatin B_1_ (EnnB_1_, ▷) and ochratoxin A (OTA, ◆) occurring in breast milk on five consecutive days from three study subjects. Detected analytes which were below the LOQ value were set to half the LOQ (LOQ/2) and the absence of displayed analytes are indicated with a star (). Other analytes were not detected in any sample.

Despite the fact that all study subjects most likely had different dietary habits, the mycotoxin occurrence pattern and concentration range were comparable. As outlined above, the diet generally is the main source of mycotoxins. The excretion into breast milk, which is considered an elimination pathway to reduce the body burden of the mother, may lead to very low chronic background exposure. However, excreted compounds are typically rather lipophilic and present in very low concentrations. For example, other environmental contaminants frequently measured and detected in breast milk include dioxins, polychlorinated biphenyls (PCBs) and perfluoroalkyl substances (PFAS). Their concentration in European breast milk was reported up to 376 ng/L and 114 ng/kg for perfluorooctanoic acid (PFOS) and WHO toxicity equivalent levels for dioxin-like compounds, respectively (Anderson *et al*. 2019; Antignac *et al*. 2016; Cariou *et al*. 2015; van den Berg *et al*. 2017). In contrast to these reports, our experiments highlight the low abundance of mycotoxins in breast milk (Braun *et al*. 2018; Braun *et al*. 2020).

In addition to breast milk, urine samples were analyzed in this smaller study. Individual results are reported in Table S7. Overall, eight mycotoxins, namely AME, CIT, DON, OTA, ZEN, α-ZAL, α-ZEL and β-ZEL were detected in these 15 urinary samples. The other 20 mycotoxins analyzed (Table S2) were not present in any sample. DON was quantified in all samples ranging from about 4,000 ng/L to 88,000 ng/L, a similar range as reported for the general Austrian population (Warth *et al*. 2012). AME and OTA were detected in 80% and 67% of all urine samples with maximum concentrations of 132 ng/L and 72 ng/L, respectively. In addition, the mycoestrogen ZEN and some of its metabolites were found in 13-67%. Here, ZEN was quantified with a maximum concentration of 642 ng/L. The occurrence of ZEN (LOD: 65 ng/L) and its estrogenic key metabolites in urine but not in breast milk (LOD of ZEN: 16 ng/L) has to be highlighted and validated in large-scale biomonitoring studies. Since these toxins are known for their capability to bind to the estrogen receptor, the excretion via maternal urine would lower the exposure of the infant. However, in a pooled breast milk sample which was obtained from a milk bank in Austria, trace amounts of ZEN (<LOQ) were found recently (Braun *et al*. 2020).

The mycotoxins AME and OTA occurred in both, breast milk and urine samples, hence, spearman correlation was conducted to evaluate their excretion pattern. No correlation was found for AME, however, the excretion via breast milk and urine was modestly associated for OTA (r=0.64, p=0.034, Figure 3C). This association might depend on the high plasma albumin binding and thus suggests a steady release of OTA to breast milk and urine (Munoz *et al*. 2014). Clearly, to establish a profound association and conclusion, a larger study collective has to be examined.

### Exposure assessment

The exposure of the infant in study A was calculated using time intervals to account for the differences in milk consumption and increase in body weight. The infants’ body weight was compared to weight-for-age standards published by the WHO (WHO 2006). Here, the initial weight after delivery and six weeks later matched the published mean weight-for-age values. In the time period of four to six months the infants’ body weight was slightly reduced compared to the mean value. Milk consumption was comparable to previous studies estimating the milk volume in the first months of life (da Costa *et al*. 2010; Kent *et al*. 1999; Neville *et al*. 1988; Salazar *et al*. 2000). The mycotoxin exposure estimates in breast milk were gained using an upper bound deterministic approach and are reported in Table 2.

**Table 2.** Estimated upper bound exposure to mycotoxins of infants based on the longitudinal occurrence data (Table S2), infants’ body weight and intake rate of breast milk in the respective time interval (Table S1).

| Interval | Days postpartum | Alternariol monomethyl ether (ng/kg bw/d) | Beauvericin | Enniatin A | Enniatin A<sub>1</sub> | Enniatin B | Enniatin B<sub>1</sub> | Ochratoxin A |
| --- | --- | --- | --- | --- | --- | --- | --- | --- |
| 1 | 0-9 | 0.12 | 0.03 | 0.01 | 0.01 | 0.06 | 0.01 | 0.04 |
| 2 | 10-19 | 0.15 | 0.09 | 0.02 | 0.03 | 0.17 | 0.03 | 0.14 |
| 3 | 20-29 | 0.95 | 0.14 | 0.03 | 0.05 | 0.22 | 0.05 | 0.25 |
| 4 | 30-39 | 0.35 | 0.19 | 0.04 | 0.06 | 0.31 | 0.05 | 0.31 |
| 5 | 40-89<sup>a</sup> | 0.46 | 0.20 | 0.04 | 0.08 | 0.31 | 0.05 | 0.38 |
| 6 | 112-183<sup>a</sup> | 0.57 | 0.20 | 0.04 | 0.08 | 0.37 | 0.07 | 0.22 |
| 7 | 211<sup>a</sup> | 0.27 | 0.23 | 0.04 | 0.07 | 0.28 | 0.08 | 0.12 |

<sup>a</sup> No samples were collected between day 89 to 112 and 183 to 211 postpartum, respectively.

Overall, the estimated exposure towards mycotoxins was very low and the highest EDI value was found for AME with 0.95 ng/kg bw per day in interval 3. However, this value can be reasonably explained and attributed to the AME peak concentration found at day 27 postpartum. AME exposure estimates in other evaluated intervals were even lower, although in most time intervals the highest estimated intake was determined for AME. For the estimates of BEA, EnnA, EnnA_1_, EnnB and EnnB_1_ an exposure trend was observed. Here, the average estimated intake increased during the first two months of life. Similarly, an increase in OTA exposure was observed up to 0.38 ng/kg bw per day, followed by a decrease to 0.12 ng/kg bw per day in the subsequent intervals. Most of these findings and the increase in exposure during the first two months can be explained by the increasing milk consumption, while the body weight and the mycotoxin concentration in breast milk were rather stable overtime.

Since it is known that infants have a less developed immune system, and limited metabolic capacities during development in the first months and years of life, this group is more susceptible towards toxic effects in general. Consequently, the European Food Safety Authority (EFSA) concluded to lower tolerable daily intake (TDI) values of substances present in food intended for infants within the first four months of life to account for the developing infant (EFSA and Hardy 2017). In addition, the EC regulation 1881/2006 specifically addresses food intended for infant consumption with MTLs being significantly lower compared to foodstuff intended for adults (EC 2006).

Of the detected toxins in the present study, only OTA is regulated in food and a tolerable weekly intake (TWI) of 120 ng/kg bw was derived by EFSA (EFSA 2010). The maximum estimated upper bound exposure during the longitudinal investigation was 0.52 ng OTA/kg bw at day 126 postpartum. Here, the OTA exposure did not exceed the hypothetically infant corrected TWI according to the recommendation by EFSA (33 ng/kg bw) or a conservative TWI of 21 ng/kg bw as derived by Kuiper-Goodman *et al*. (2010). Although, the infant corrected TWI values were not exceeded in the studied time frame, potential uncertainties, e.g. resulting from the conducted sample pooling strategy, remain and have to be addressed in the future.

In comparison to our estimates, others who derived OTA exposure for Chilean infants (Munoz *et al*. 2014) reported nearly 100-fold higher EDI values with average values between 5 to 12 ng/kg bw per day. However, worldwide reported occurrence data of breast milk OTA levels vary considerably and likely explain these differences in EDI values. As discussed above, complex lactational transfer processes, changes in breast milk composition, demand of the infant and mammary gland physiology are key factors for the transmission of substances to breast milk (LaKind *et al*. 2009).

Comparing our preliminary results with the estimates derived from Nigerian breast milk (Braun *et al*. 2018), the EDI values of BEA, EnnB and OTA were around 12x, 6x and 30x lower in this study, respectively. The main reason for this divergence is the maximum concentration of mycotoxins measured which were, as one would estimate, slightly higher for all three toxins in Nigerian breast milk.

The mycoestrogen ZEN was not detected in any sample in this pilot study, and recently detected only below the LOQ value in one pooled Austrian breast milk sample obtained from a milk bank (Braun *et al*. 2018; Braun *et al*. 2020). These data suggest that most reports overestimate infant’s exposure to mycotoxins. For example, Massart *et al*. (2016) reported a mean ZEN concentration of 1130 ng/L, which may be the result of using less specific analytical approaches. Here, cross-reactivity reactions (ELISA) or interfering peaks (LC-FD) may be interpreted as positive signals. In contrast, LC-MS/MS approaches clearly benefit by using stable isotope reference standards to minimize the probability of reporting false-positives. Thus, our pilot study revealed that overall only the most sensitive and rather lipophilic analytes were identified in very low concentration levels. Still, low-dose combinatory effects of mycotoxins have to be evaluated in detail as combinations of mycotoxins have shown to exert synergistic or antagonistic effects *in vitro* (Vejdovszky *et al*. 2017; Vejdovszky *et al*. 2016). In our experiments, these mycotoxins were either not detected (ZEN, α-ZEL and β-ZEL) or were not extractable with the utilized method (AOH). However, exposure in early developmental stages may be possible, as it is known that e.g. ZEN and its metabolites can be transferred across the human placental barrier (Warth *et al*. 2019).

Conjugated mycotoxins, such as glucuronidated or sulfated forms, were not assessed in the breast milk samples, as the analytical method was not validated for such an approach. In an earlier study, enzymatic treatment was applied, and no increasing mycotoxin concentrations were observed (Braun *et al*. 2018). However, the potential presence of conjugated forms might increase the overall exposure of the parent mycotoxins if these are released in the intestine of infants. In addition, infants might be exposed to other compounds such as xenoestrogens as shown recently (Preindl *et al*. 2019). In light of the new exposome paradigm, low-dose mixture effects of mycotoxins with persistent organic pollutants like dioxins or PFOS may be relevant and need to be addressed in the future to evaluate their combined effects in a holistic manner.

However, replacing breast milk by feeding alternatives like infant formula may lead to an increased risk for the infant. We could recently demonstrate that infants are exposed to mycotoxins via complementary infant food to a higher extent than via breast milk (Braun et al. unpublished). Consequently, potential low mycotoxin exposure via breast milk should clearly not be a factor to reduce or avoid breastfeeding.

### Study design: Strengths, limitations and future perspectives

The main strength of the study is the assessment of longitudinal exposure data (study A) of a multitude of mycotoxins and key metabolites in this important bio-fluid. These data were used to calculate the exposure of the infant by taking actually measured milk consumption and body weight into account. In addition, the results of study A could be verified by proof-of-principle experiments in a shorter time frame (study B). Based on these data it was possible to assess inter-day and inter-individual mycotoxin occurrence patterns. The main limitation of the presented data is that the sample preparation procedure was not suitable to extract certain toxins (e.g. the polar trichothecenes DON, NIV, the fumonisins, AOH, and the AFB_1_-N7-guanine adduct) as described before (Braun *et al*. 2020). However, these are generally more polar and considering their rapid excretion via urine not very likely to be transferred to breast milk without a suitable carrier. Another important limitation is the low number of participants who donated breast milk to draw robust conclusions from the generated data set and extrapolate to a general population.

However, the preliminary results of study A and B resulted in new insights in an under-researched field of early-life exposure assessment. To better suit epidemiological approaches in the future, the study design might need some adjustments. While highly valuable, longitudinal assessments may not be feasible in large-scale human biomonitoring (HBM) studies considering the intensive sampling efforts and the required dedication of mothers during a time frame over several months.

In study A and B, the coefficient of variation (CV) for mycotoxins detected in more than 80% of analyzed samples were below 54%, except for AME with CVs above 90%. In study A, only minor differences in CVs were observed by comparing the variability in occurrence of each mycotoxin per defined time interval. Hence, future HBM studies should aim to include short time intervals of around 10-15 days to account for changes in intra-individual occurrence and variability as well as differences in the metabolism and excretion of mycotoxins. In addition, several samples collected throughout the day might be pooled to obtain representative samples. Although this simplified sample collection strategy might be suitable, differences in fore- and hindmilk pose a potential source of variability. For example, foremilk is rich in proteins, whereas hindmilk has higher lipid concentrations, *i.e* the latter might result in the detection of more lipophilic analytes. Especially in exposome-scale studies, this approach would introduce a higher uncertainty factor or analytes of potential interest might be missed if only hindmilk is used. These differences should be addressed by comparing foremilk, hindmilk and a combination thereof in follow-up experiments.

## Conclusion and outlook

In this paper we report the first comprehensive longitudinal investigation of mycotoxin mixtures in breast milk (study A). Our findings indicated that mycotoxin concentrations in the analyzed Austrian breast milk samples were low. However, extrapolation of these results for the general population were not possible and have to be addressed in studies including more study subjects in the future. In study A, regulated mycotoxins such as aflatoxin, ZEN or their key metabolites were not detected in any sample. Only rather lipophilic and most sensitive analytes were found in very low concentration ranges suggesting chronic background exposures through the diet. In light of the preliminary data gained in study B, inter- and intraindividual effects were assessed and demonstrated to be minimal. Thus, sampling in time intervals of 10 to 15 days was deemed to be sufficient to retrieve exposure data for the detected toxins.

Based on the insights gained in this pilot study, the exposure of the Austrian infant to mycotoxins via breast milk was estimated to be negligible. When comparing previous EDI values to our calculations, we conclude that most published data overestimated the exposure of infants to mycotoxins. However, factors influencing contamination patterns in breast milk have to be further investigated using large-scale HBM studies to assess intra-, inter-individual and geographical aspects. In addition, our findings of low chronic background exposures with mostly non-regulated toxins have to be verified in large-scale human biomonitoring (HBM) studies. These should aim to generate exposome-scale data by non-targeted mass spectrometry-based approaches, to better understand relevant factors and assess combinatory effects of e.g. mycoestrogens. Overall, the potential low-level presence of mycotoxins clearly does not warrant to discourage mothers from breastfeeding.

## Declaration of interest

None.

## Acknowledgements

The authors want to gratefully thank all participants who provided samples. We would like to acknowledge Ipek Savaş and Patrick Windisch for skillful practical assistance. In addition, we want to acknowledge the Mass Spectrometry Centre (MSC) of the Faculty of Chemistry at the University of Vienna and Sciex for providing mass spectrometric instrumentation. This work was performed with the financial support of the University of Vienna and the Austrian Science Fund (FWF): P 33188-B.

## Appendix: Supplementary material

Additional information is available on mycotoxin extraction from urine, LC-MS/MS analysis and in-house validation for urine samples, analytical method performance for breast milk measurements (Table S1, MS and MS/MS parameters used for urinary mycotoxin detection (Table S2), analytical method performance in urine (Table S3), infant’s body weight and milk consumption (Table S4), individual sample concentration levels of mycotoxins found in breast milk in study A (Table S5) and study B (Table S6), as well as concentration levels of mycotoxins found in individual urine samples in study B (Table S7).

## Notes

### Competing Interest Statement

The authors have declared no competing interest.

### Summary of Updates

Overall revision of the manuscript and urinary mycotoxin occurrence data were added in the smaller study B.

