## Supplementary Material for "Longitudinal assessment of mycotoxin co-exposures in exclusively breastfed infants"

### 1. Supplementary Materials and Methods:

#### 1.1. Urinary sample preparation and LC-MS/MS analysis

Mycotoxins were extracted from urine and analyzed by LC-MS/MS following the protocol of Šarkanj *et al.* (2018) with minor modifications and an expanded set of analytes. In brief, samples were thawed, homogenized and a 1 mL aliquot was centrifuged (3,000 x g, 2 min, 22 °C). An aliquot of the supernatant (500 µL) was incubated for 16 h at 37 °C with phosphate-buffered saline (PBS, 500 µL, 200 mM, pH = 7.4), which contained β-glucuronidase from *E.coli* (3,000 U; Type IX-A, Sigma-Aldrich). Then, 1 mL crude sample extract was loaded onto a pre-equilibrated (1 mL MeOH and 1 mL $H_2O$) Oasis HLB PRiME solid phase extraction (SPE) column (1cc, 30 mg, Waters, Milford, MA). The SPE column was washed using 500 µL $H_2O$ twice, while analytes of interest were eluted in three steps using each time 200 µL ACN. This extract was subsequently evaporated to dryness at 4 °C using a vacuum concentrator (Labconco, Missouri, USA) and samples were reconstituted in 490 µL ACN/$H_2O$/acetic acid (10/89.9/0.1, v/v/v) and 10 µL of the internal standard mixture (IS-mix) was added. After homogenization the samples were transferred to amber LC-vials and analyzed.

The LC-MS/MS system used for mycotoxin quantification in urine was the same used for breast milk analysis. Chromatographic separation of mycotoxins was achieved using an Acquity HSS T3 column (1.8 µm, 2.1x100 mm) guarded with a VanGuard pre-column (1.8 µm, Waters, Vienna, Austria) at a flow rate of 0.1 mL/min. A binary gradient elution was used consisting of eluent A ($H_2O$) and B (ACN), both acidified with 0.1% acetic acid. During the first two minutes, the column was kept at 10% eluent B, while the percentage of B was linearly raised in the next 13 min to 50% and within 3 min to 95%. The gradient was held for 4 min at 95% B and subsequently, using a rapid decrease to initial conditions within 0.1 min, the column was re-equilibrated at 10% B. Overall, this resulted in a chromatographical runtime of 25 min. The instruments' autosampler and column oven were maintained at 7 °C and 35 °C, respectively. For analysis, a sample volume of 10 µL was injected onto the column. The MS was acquiring in scheduled multiple reaction monitoring (sMRM) applying fast polarity switching using a detection window of 180 seconds around the expected retention time of each analyte (Table S2).

Analytes were optimized using the software's integrated optimization tool via direct infusion of reference standards. Thus, preferred MRM parameters as described in Table S2 were determined for all analytes. The ESI ion source was operated with following parameters: source temperature 450°C, curtain gas 30 psi, collision gas high, ion source gases (sheath and drying gas) 60 and 40 psi and the ion spray voltage was set to 4500 V in positive and -4500 V in negative mode.

##### 1.2. Validation experiments in urine

In house validation was performed according to established guidelines, e.g. the European Commission Decision 2002/657/EC (EC 2002) and the EuraChem Laboratory Guide (Magnusson 2014). The following parameters were evaluated in-house: linearity, extraction recovery ($R_E$), repreatability ($RSD_r$), intermediate precision ($RSD_R$), selectivity, specificity, limit of detection (LOD), limit of quantification (LOQ) and signal suppression or enhancement (SSE). Calibration standards were prepared in neat solvent and unspiked pooled urine extracts (matrix-matched calibrants). Matrix-matched calibration curve (1/x weighted) was established using at least five concentration levels per mycotoxin. The standard addition method was used in cases where the unspiked urine extract was naturally contaminated. Peak area ratios were used for quantification for mycotoxins where internal standards were available, while all other mycotoxins were quantified using the peak area instead. LOD and LOQ values were calculated by evaluating the signal to noise level. A signal to noise ratio of 3:1 and 6:1 from spiked urine chromatograms were set to the LOD and LOQ value, respectively.

#### 2. Supplementary Results and Discussion

##### 2.1. In-house validation – biomarkers of mycotoxin exposure in urine

The method performance was validated in-house according to established guidelines (EC 2002; Magnusson 2014). Since, no certified reference material was available, a non-spiked pooled urine sample, obtained from one individual who tried to avoid mycotoxin prone diet, was used for all validation experiments. The following parameters were evaluated: sensitivity, selectivity, linearity, repeatability, intermediate precision, extraction recovery and matrix effects. Overall, method validation was successful, and the detailed results are reported in Table S3.

The method allows the quantification of 28 of the 30 selected mycotoxins, mostly in the lower ng/mL range. LOD and LOQ values ranged from 0.005 ng/mL to 4.6 ng/mL and 0.01 ng/mL to 9.2 ng/mL, respectively. Low LOD values were achieved for most major aflatoxins ($AFB_1$, $AFB_2$, $AFG_1$, $AFM_1$), ochratoxin A (OTA) and sterigmatocystin, which could all be detected below 0.045 ng/mL. LOD values of other mycotoxins were <0.9 ng/mL, except $AFQ_1$ and T-2 toxin with LODs of 1.8 ng/mL and 4.6 ng/mL, respectively. Sensitivity was lower compared to a previously validated method in urine (Šarkanj *et al.* 2018). However, this difference can be mostly explained by the addition of 18 more analytes which eluted in a time window close to those analytes which were included in the previous method. Hence, dwell times during the MS/MS experiments were reduced, lowering the overall signal-to-noise ratio. The method's selectivity was evaluated by comparing non-spiked with spiked samples. If no co-eluting peak with a signal to noise ratio >3 was found, proper detection was considered feasible. Overall, identification in matrix-matched samples was evaluated based on following criteria:

retention time, quantifier and qualifier ion, as well as the ion ratio. Linearity of each compound was deemed acceptable, if a regression coefficient of $R^2 > 0.99$ was obtained using matrix-matched calibrants. Extraction recoveries as stated in Table S3 were in agreement with established guidelines, except for $AFB_1$-N7-guanine adduct, $AFG_2$, alternariol, citrinin, and fumonisin $B_1$ ($FB_1$). These were either not sufficiently extracted, or, in case of $AFG_2$ and AOH, interfering matrix peaks were observed in all evaluated MRM transitions, which hindered proper evaluation. $AFQ_1$, ochratoxin B, ochratoxin alpha, sterigmatocystin and T-2 toxin passed the validation criteria on the higher spiking level, however, the lower spiking level was not reported, since the spiked concentration was lower than the finally assigned LOQ value. However, extraction recoveries and variability for these analytes were in the range of the higher spiking level and thus, the method can be used as a screening tool if these analytes are close to their evaluated LOQ values.

Overall, $RSD_r$ and $RSD_R$ of successfully validated mycotoxins were in good agreement with established guidelines (EC 2002). Here, variability ($RSD_r$ and $RSD_R$) on both evaluated spiking levels was below 21%, except for OTA in the higher spiking experiment ($RSD_R$ of 36%). However, this slightly higher variability of OTA as well as the extraction efficiency of OTA and $FB_1$ can be compensated by adding the IS-mix prior to any extraction or clean-up step in the future application of this method in epidemiological studies. SSE was assessed by comparing calibration slopes of matrix-matched calibrant to neat solvent calibrants and are reported as average values. Overall, SSE was minor with 80-101% for most aflatoxins, whereas highest signal suppression was observed for alternariol monomethyl ether and the group of zearalenone, zearalenone and their key metabolites with SSEs of 26-56%. Zearalenone (SSE of 101%), and other mycotoxins where IS were available, was evaluated and corrected using the peak area ratio of $^{12}C/^{13}C$ and matrix effects can thus be neglected.

Overall, the method performance proved to be fit for purpose to detect and quantify mycotoxins in urine. This multi-analyte method required a generic solid phase extraction clean-up protocol, hence, some compromises in the methods performance were tolerated.

#### 3. Supplementary Tables:

**Table S1.** Performance characteristics of the method for the extraction of mycotoxins in breast milk as obtained during in-house validation on the QTrap6500$^+$ instrument including regression coefficient ($R^2$), spiking level, recovery of the extraction step ($R_E$), intermediate precision ($RSD_R$), repeatability ($RSD_r$), limits of detection (LOD) and limits of quantification (LOQ). Modified from Braun *et al.* (2020).

| Analyte | Regression coefficients | Spiking level | $R_E \pm RSD_R$ | $RSD_r$[b] | LOD | LOQ |
|---|---|---|---|---|---|---|
| | $R^2$ | [ng/L] | [%] | [%] | [ng/L] | [ng/L] |
| Aflatoxicol | 0.999 | 50 | 91 ± 3 | 3 | 30 | 60 |
| Aflatoxin $B_1$ | 0.999 | 10 | 96 ± 2 | 2 | 2.5 | 5.0 |
| Aflatoxin $B_2$ | 0.999 | 20 | 100 ± 5 | 6 | 1.0 | 2.0 |
| Aflatoxin $G_1$ | 0.999 | 30 | 99 ± 1 | 2 | 3.5 | 7.0 |
| Aflatoxin $G_2$ | 0.999 | 30 | 100 ± 8 | 9 | 4.0 | 8.0 |
| Aflatoxin $M_1$ | 0.999 | 10 | 109 ± 4 | 5 | 2.0 | 4.0 |
| Aflatoxin $M_2$ | 0.999 | 48 | 87 ± 13 | 20 | 14 | 28 |
| Aflatoxin $P_1$ | 0.999 | 48 | 91 ± 7 | 6 | 9.0 | 18 |
| Aflatoxin $Q_1$ | 0.999 | 10 | 89 ± 8 | 6 | 13 | 26 |
| Aflatoxin $B_1$-N7-guanine | 0.999 | 20 | 39 ± 19 | 26 | 4.0 | 8.0 |
| Alternariol[a] | 0.999 | 50 | - | - | 4.0 | 8.0 |
| Alternariol monomethyl ether | 0.999 | 10 | 86 ± 5 | 5 | 0.5 | 1.0 |
| Beauvericin | 0.999 | 6 | 85 ± 9 | 10 | 0.1 | 0.2 |
| Citrinin | 0.999 | 6 | 118 ± 18 | 15 | 3.0 | 6.0 |
| Deoxynivalenol[a] | 0.999 | 720 | - | - | 106 | 212 |
| Dihydrocitrinone | 0.999 | 96 | 55 ± 7 | 4 | 14 | 28 |
| Enniatin A | 0.999 | 6 | 99 ± 5 | 5 | 0.5 | 1.0 |
| Enniatin $A_1$ | 0.999 | 6 | 98 ± 5 | 4 | 0.9 | 1.8 |
| Enniatin B | 0.999 | 6 | 85 ± 6 | 9 | 0.7 | 1.4 |
| Enniatin $B_1$ | 0.999 | 6 | 95 ± 6 | 4 | 0.5 | 1.0 |
| HT-2 toxin | 0.996 | 720 | 84 ± 12 | 9 | 300 | 600 |
| Nivalenol[a] | 0.999 | 800 | - | - | 70 | 140 |
| Ochratoxin A | 0.999 | 30 | 96 ± 3 | 4 | 0.8 | 1.5 |
| Ochratoxin B | 0.999 | 20 | 97 ± 3 | 3 | 2.5 | 5.0 |
| Ochratoxin α | 0.998 | 160 | 83 ± 18 | 24 | 24 | 48 |
| Sterigmatocystin | 0.999 | 15 | 90 ± 2 | 2 | 0.5 | 1.0 |
| T-2 toxin | 0.999 | 96 | 106 ± 5 | 6 | 11 | 22 |
| Tentoxin | 0.999 | 96 | 101 ± 4 | 5 | 23 | 46 |
| Zearalanone | 0.999 | 96 | 92 ± 4 | 3 | 60 | 120 |
| α-Zearalanol | 0.999 | 128 | 103 ± 3 | 3 | 73 | 146 |
| β-Zearalanol | 0.999 | 128 | 98 ± 4 | 5 | 75 | 150 |
| Zearalenone | 0.999 | 96 | 103 ± 5 | 4 | 16 | 32 |
| α-Zearalenol | 0.999 | 100 | 90 ± 5 | 5 | 44 | 87 |
| β-Zearalenol | 0.999 | 100 | 92 ± 5 | 5 | 54 | 108 |

[a] AOH, DON and NIV could not be recovered following our extraction procedure. Therefore, these three toxins were not considered to be successfully validated.

**Table S2.** MS and MS/MS parameters of the method used for the determination of 30 mycotoxins/mycotoxin metabolites in urine, including retention time (RT), mass-to-charge-ratios (*m/z*) of precursor ion and product ions, declustering potential (DP), collision energy (CE), collision cell exit potential (CXP) and the ion ratio.

| Analyte | RT | Precursor ion | Ion species | Product ion (Quantifier/qualifier) | DP | CE | CXP | Ion ratio[a] |
|---|---|---|---|---|---|---|---|---|
| | min | *m/z* | | *m/z* | V | V | V | % |
| Aflatoxicol | 18.1 | 297.0 | $[M-H_2O+H]^+$ | 269.1/115.1 | 71 | 29/83 | 12/14 | 0.81 |
| Aflatoxin $B_1$ | 17.6 | 313.0 | $[M+H]^+$ | 241.0/213.0 | 106 | 49/61 | 14/16 | 0.66 |
| Aflatoxin $B_2$ | 16.7 | 315.0 | $[M+H]^+$ | 243.0/203.0 | 125 | 53/49 | 16/12 | 0.52 |
| Aflatoxin $G_1$ | 16.7 | 329.1 | $[M+H]^+$ | 200.0/243.1 | 86 | 59/39 | 12/14 | 1.61 |
| Aflatoxin $G_2$ | 15.8 | 331.1 | $[M+H]^+$ | 313.2/245.2 | 111 | 35/43 | 18/14 | 0.54 |
| Aflatoxin $M_1$ | 14.7 | 329.1 | $[M+H]^+$ | 273.2/229.1 | 91 | 35/59 | 16/12 | 0.55 |
| $^{13}C$-Aflatoxin $M_1$ | 14.7 | 346.0 | $[M+H]^+$ | 288.2 | 91 | 35 | 16 | - |
| Aflatoxin $M_2$ | 13.8 | 331.0 | $[M+H]^+$ | 259.0/241.0 | 96 | 33/57 | 16/14 | 0.59 |
| Aflatoxin $P_1$ | 14.6 | 299.1 | $[M+H]^+$ | 270.7/215.1 | 126 | 35/38 | 18/11 | 0.32 |
| Aflatoxin $Q_1$ | 14.9 | 328.7 | $[M+H]^+$ | 206.0/177.0 | 121 | 33/47 | 14/12 | 0.69 |
| Aflatoxin-N7-guanine | 11.0 | 480.0 | $[M+H]^+$ | 152.1/135.0 | 46 | 23/60 | 10/14 | 0.15 |
| Alternariol | 18.4 | 257.0 | $[M-H]^-$ | 215.0/212.1 | -110 | -34/-38 | -13/-12 | 1.66 |
| $^{2}H_4$-Alternariol | 18.4 | 261.0 | $[M-H]^-$ | 150.0 | -110 | -46 | -5 | - |
| Alternariol monomethyl ether | 20.7 | 271.1 | $[M-H]^-$ | 256.0/227.0 | -95 | -32/-50 | -13/-9 | 0.17 |
| Citrinin | 19.0 | 251.0 | $[M+H]^+$ | 233.1/205.1 | 36 | 23/37 | 14/18 | 0.11 |
| $^{13}C$-Citrinin | 19.0 | 264.0 | $[M+H]^+$ | 217.2 | 56 | 37 | 10 | - |
| Dihydrocitrinone | 16.0 | 265.0 | $[M-H]^-$ | 176.9/246.9 | -30/-45 | -34/-26 | -13/-15 | 0.70 |
| Deoxynivalenol | 8.5 | 354.9 | $[M+CH_3COO]^-$ | 265.0/59.0 | -30 | -20/-58 | -21/-27 | 2.94 |
| $^{13}C$-Deoxynivalenol | 8.5 | 370.1 | $[M+CH_3COO]^-$ | 278.8 | -20 | -22 | -15 | - |
| Deepoxy-deoxynivalenol | 10.2 | 280.9 | $[M+H]^+$ | 109.1/137.1 | 36 | 21/19 | 12/10 | 0.96 |
| Nivalenol | 5.7 | 371.0 | $[M+CH_3COO]^-$ | 281.1/59.1/203.0 | -20 | -20/-40/-30 | -19/-9/-13 | 0.86 |
| $^{13}C$-Nivalenol | 5.7 | 386.0 | $[M+CH_3COO]^-$ | 295.0 | -75 | -22 | -15 | - |
| Fumonisin $B_1$ | 14.9 | 722.5 | $[M+H]^+$ | 334.4/352.3 | 121 | 57/55 | 4/12 | 0.98 |
| $^{13}C$-Fumonisin $B_1$ | 14.9 | 756.3 | $[M+H]^+$ | 356.3 | 130 | 46 | 10 | - |
| Ochratoxin A | 20.6 | 404.0 | $[M+H]^+$ | 239.0/102.0 | 91 | 37/105 | 16/14 | 0.33 |
| $^{13}C$-Ochratoxin A | 20.6 | 424.0 | $[M+H]^+$ | 250.0 | 51 | 33 | 12 | - |
| Ochratoxin B | 19.5 | 370.1 | $[M+H]^+$ | 205.0/103.1 | 86 | 33/77 | 12/16 | 0.32 |
| Ochratoxin α | 15.0 | 254.9 | $[M-H]^-$ | 166.9/123.0/110.9 | -90 | -36/-40/-44 | -11/-17/-21 | 0.16 |
| Sterigmatocystin | 21.1 | 325.1 | $[M+H]^+$ | 281.1/310.2 | 96 | 51/35 | 16/18 | 0.85 |
| T-2 toxin | 20.3 | 467.3 | $[M+H]^+$ | 215.2/185.1 | 56 | 29/31 | 18/11 | 0.78 |
| Tentoxin | 18.2 | 413.3 | $[M-H]^-$ | 141.0/271.1 | -105 | -30/-24 | -11/-15 | 0.56 |
| Zearalanone | 20.7 | 319.1 | $[M-H]^-$ | 107.0/161.0 | -125 | -40/-38 | -13/-15 | 1.32 |
| α-Zearalanol | 19.6 | 321.1 | $[M-H]^-$ | 277.1/161.0 | -120 | -30/-38 | -18/-9 | 0.08 |
| β-Zearalanol | 18.8 | 321.2 | $[M-H]^-$ | 277.1/303.1 | -120 | -30/-30 | -18/-20 | 0.27 |
| Zearalenone | 20.8 | 317.1 | $[M-H]^-$ | 131.0/175.0 | -110 | -42/-34 | -8/-13 | 1.20 |
| $^{13}C$-Zearalenone | 20.8 | 335.2 | $[M-H]^-$ | 185.1 | -110 | -34 | -13 | - |
| α-Zearalenol | 19.8 | 319.2 | $[M-H]^-$ | 160.0/130.1 | -115 | -44/-50 | -13/-20 | 0.63 |
| β-Zearalenol | 19.0 | 319.1 | $[M-H]^-$ | 160.0/130.1 | -115 | -44/-50 | -13/-20 | 0.65 |

[a] Calculated as the ratio of qualifier and quantifier and expressed in percent.

**Table S3.** Performance characteristics of the method to quantify mycotoxins in urine as obtained during in-house validation including regression coefficient ($R^2$), spiking levels, recoveries of the extraction step ($R_E$), intermediate precision ($RSD_R$), repeatability ($RSD_r$), signal suppression/enhancement (SSE), limits of detection (LOD) and limits of quantification (LOQ).

| Analyte | $R^2$ | Spiking levels[a] | $R_E \pm RSD_R$ | | $RSD_r$ | | SSE[b] | LOD | LOQ |
|---|---|---|---|---|---|---|---|---|---|
| | | | level 1 | level 2 | level 1 | level 2 | | | |
| | | (ng/L) | (%) | (%) | (%) | (%) | (%) | (ng/L) | (ng/L) |
| Aflatoxicol | 0.999 | 800/13333 | 87 ± 20 | 94 ± 11 | 14 | 6 | 69 | 170 | 340 |
| Aflatoxin $B_1$ | 0.999 | 120/2000 | 100 ± 18 | 97 ± 8 | 5 | 5 | 81 | 30 | 60 |
| Aflatoxin $B_2$ | 0.999 | 200/3333 | 79 ± 12 | 98 ± 8 | 10 | 6 | 92 | 45 | 90 |
| Aflatoxin $G_1$ | 0.999 | 140/2333 | 87 ± 13 | 97 ± 10 | 9 | 6 | 99 | 40 | 80 |
| Aflatoxin $G_2$[c] | - | 200/3333 | - | - | - | - | - | - | - |
| Aflatoxin $M_1$ | 0.999 | 140/2333 | 96 ± 19 | 97 ± 6 | 2 | 4 | 97 | 30 | 60 |
| Aflatoxin $M_2$ | 0.999 | 600/10000 | 84 ± 11 | 97 ± 9 | 3 | 5 | 81 | 115 | 230 |
| Aflatoxin $P_1$ | 0.999 | 600/10000 | 74 ± 13 | 90 ± 8 | 4 | 5 | 66 | 110 | 220 |
| Aflatoxin $Q_1$[d] | 0.999 | 600/10000 | - | 79 ± 8 | - | 5 | 100 | 1800 | 3600 |
| Aflatoxin-N7-guanine[c] | 0.999 | 40/667 | - | 51 ± 32 | - | 23 | 49 | 30 | 60 |
| Alternariol[c] | - | 1000/16667 | - | - | - | - | 79 | - | - |
| Alternariol monomethyl ether | 0.999 | 60/1000 | 60 ± 14 | 66 ± 17 | 8 | 18 | 27 | 12 | 25 |
| Citrinin[e] | 0.998 | 3000/50000 | 36 ± 25 | 32 ± 53 | 8 | 34 | 68 | 700 | 1400 |
| Deepoxy-deoxynivalenol | 0.999 | 2000/30000 | 71 ± 18 | 97 ± 13 | 20 | 9 | 37 | 470 | 940 |
| Deoxynivalenol | 0.999 | 4000/66667 | 86 ± 17 | 105 ± 13 | 6 | 7 | 92 | 950 | 1900 |
| Dihydrocitrinone | 0.999 | 1200/20000 | 77 ± 14 | 92 ± 11 | 16 | 12 | 99 | 300 | 600 |
| Fumonisin $B_1$[e] | 0.999 | 400/6667 | 35 ± 17 | 15 ± 36 | 8 | 24 | 88 | 70 | 140 |
| Nivalenol | 0.999 | 1000/16667 | 103 ± 14[e] | 94 ± 8 | 5 | 7 | 85 | 300 | 600 |
| Ochratoxin A[e] | 0.998 | 40/667 | 58 ± 21 | 68 ± 36 | 20 | 52 | 96 | 20 | 40 |
| Ochratoxin B[d] | 0.998 | 40/667 | - | 73 ± 35 | - | 19 | 87 | 30 | 60 |
| Ochratoxin α[d] | 0.998 | 200/3333 | - | 90 ± 40 | - | 22 | 82 | 150 | 300 |
| Sterigmatocystin[d] | 0.999 | 10/167 | - | 77 ± 11 | - | 14 | 95 | 5 | 10 |
| T-2 toxin[d] | 0.999 | 6000/100000 | - | 92 ± 9 | - | 7 | 58 | 4600 | 9200 |
| Tentoxin | 0.999 | 200/3333 | 89 ± 7 | 93 ± 7 | 3 | 4 | 42 | 45 | 90 |
| Zearalanone | 0.999 | 600/10000 | 94 ± 14 | 103 ± 11 | 4 | 9 | 26 | 195 | 390 |
| Zearalenone | 0.999 | 400/6667 | 80 ± 18 | 93 ± 11 | 8 | 12 | 101 | 65 | 130 |
| α-Zearalanol | 0.999 | 1200/20000 | 90 ± 17 | 85 ± 18 | 20 | 14 | 37 | 265 | 530 |
| α-Zearalenol | 0.999 | 200/3333 | 86 ± 19 | 88 ± 14 | 15 | 8 | 31 | 100 | 200 |
| β-Zearalanol | 0.999 | 1200/20000 | 73 ± 18 | 98 ± 14 | 15 | 19 | 49 | 190 | 380 |
| β-Zearalenol | 0.999 | 200/3333 | 82 ± 17 | 101 ± 19 | 18 | 13 | 56 | 65 | 130 |

[a] Spiking levels are reported in the following order: level 1 / level 2.

[b] Calculated as the ratio of matrix-matched calibration slope and solvent calibration slope and expressed as percent.

[c] $AFB_1$-N7-guanine, $AFG_2$ and AOH could not be sufficiently recovered. Therefore, these three toxins were not successfully validated on any level.

[d] $AFQ_1$, OTB, OTα, STC and T-2 were not reported on spiking level 1, as the spiked concentration was below the final LOQ value. However, extraction efficiency and variability were in a similar range as spiking level 2. Thus, this method can be used as a screening method in the lower range.

[e] In following experiments, their extraction efficiency can be considered 100%, as $^{13}C$ labelled reference standard will be spiked prior to any sample clean-up procedure.

**Table S4.** Basic descriptive data for infant's body weight and breast milk intake on each measured day postpartum and the respective assigned interval.

| Interval | Days postpartum | Infant weight (kg) | Intake rate (L/d) |
|---|---|---|---|
| 1 | 0 (delivery) | 3.49 | 0.10 |
| 2 | 10 | 3.64 | 0.25 |
| 3 | 20 | 3.80 | 0.40 |
| 4 | 30 | 3.95 | 0.55 |
| 5 | 40 | 4.18 | 0.70 |
| 6 | 112 | 5.30 | 0.90 |
| 7 | 211 | 6.07 | 1.0 |

**Table S5.** Concentration levels of mycotoxins found in breast milk samples (n=87) obtained with study A during the first 211 days postpartum.

| Analyte | AME | BEA | EnnA | EnnA$_1$ | EnnB | EnnB$_1$ | OTA |
|---|---|---|---|---|---|---|---|
| **Retention time (min)** | 8.1 | 11.0 | 11.5 | 11.3 | 10.8 | 11.1 | 6.4 |
| **LOD (ng/L)[a]** | 0.5 | 0.1 | 0.5 | 0.9 | 0.65 | 0.5 | 0.8 |
| **LOQ (ng/L)[a]** | 1.0 | 0.3 | 1.0 | 1.8 | 1.3 | 1.0 | 1.5 |
| **Regression coefficient $R^2$** | 0.999 | 0.999 | 0.999 | 0.999 | 0.999 | 0.999 | 0.999 |

| Sample day postpartum | AME (ng/L) | BEA | EnnA | EnnA$_1$ | EnnB | EnnB$_1$ | OTA |
|---|---|---|---|---|---|---|---|
| Delivery | n.a[b] | n.a | n.a | n.a | n.a | n.a | n.a |
| 3 | 3.0 | 0.8 | <LOD[c] | <LOD | <LOQ | <LOD | 2.0 |
| 4 | 6.0 | 1.1 | <LOD | <LOD | 1.6 | <LOD | 1.8 |
| 5 | 1.8 | 1.2 | <LOD | <LOD | 1.3 | <LOD | 2.2 |
| 6 | 2.0 | 1.3 | <LOD | <LOD | 2.0 | <LOQ[d] | <LOQ |
| 7 | 11 | 1.1 | <LOD | <LOD | 2.8 | <LOQ | <LOQ |
| 8 | 3.9 | 1.3 | <LOD | <LOD | 3.2 | <LOQ | 1.6 |
| 9 | 2.4 | 1.3 | <LOD | <LOD | 2.8 | <LOD | <LOQ |
| 10 | 1.9 | 1.3 | <LOQ | <LOD | 3.2 | <LOQ | 2.5 |
| 11 | 1.7 | 1.3 | <LOD | <LOD | 2.4 | <LOQ | 2.1 |
| 12 | 2.2 | 1.2 | <LOD | <LOD | 1.5 | <LOD | 1.9 |
| 13 | 1.9 | 1.2 | <LOQ | <LOD | 3.3 | <LOQ | 1.9 |
| 14 | 4.6 | 1.1 | <LOD | <LOD | 2.1 | <LOD | 1.9 |
| 16 | <LOD | 1.4 | <LOD | <LOD | 2.4 | <LOQ | 2.0 |
| 15 | 2.2 | 1.3 | <LOD | <LOD | 2.9 | <LOQ | 1.9 |
| 17 | 2.3 | 1.5 | <LOD | <LOD | 2.5 | <LOQ | 2.3 |
| 18 | 3.1 | 1.3 | <LOD | <LOD | 2.3 | <LOD | 1.7 |
| 19 | 1.9 | 1.1 | <LOD | <LOD | 2.2 | <LOD | 2.1 |
| 20 | 2.1 | 1.6 | <LOD | <LOD | 2.6 | <LOQ | 2.1 |
| 21 | 2.1 | 1.4 | <LOD | <LOD | 2.1 | <LOQ | 2.0 |
| 22 | 2.1 | 1.3 | <LOQ | <LOD | 2.9 | <LOQ | 3.0 |
| 23 | 1.9 | 1.3 | <LOD | <LOD | 2.5 | <LOQ | 2.2 |
| 25 | 2.5 | 1.0 | <LOD | <LOD | 1.4 | <LOQ | 2.7 |
| 26 | 2.0 | 1.1 | <LOD | <LOD | <LOQ | <LOD | 2.6 |
| 27 | 58 | 1.4 | <LOD | <LOD | 1.4 | <LOQ | 2.2 |
| 29 | 2.0 | 1.4 | <LOD | <LOD | 3.6 | <LOQ | 2.0 |
| 31 | 2.0 | 1.1 | <LOD | <LOD | 2.1 | <LOD | 1.8 |
| 32 | 1.8 | 1.5 | <LOD | <LOD | 2.1 | <LOQ | 1.7 |
| 33 | 2.4 | 1.3 | <LOD | <LOD | 1.7 | <LOD | <LOQ |
| 34 | 3.6 | 1.2 | <LOQ | <LOD | 3.0 | <LOD | 1.9 |
| 35 | 3.1 | 1.3 | <LOD | <LOD | 2.6 | <LOQ | 3.0 |
| 37 | 2.0 | 1.3 | <LOD | <LOD | 2.7 | <LOQ | 2.9 |
| 38 | 1.8 | 1.3 | <LOD | <LOD | 2.0 | <LOQ | 3.0 |
| 39 | 2.3 | 1.2 | <LOD | <LOD | 2.0 | <LOD | 2.9 |
| 40 | 3.7 | 1.7 | <LOD | <LOD | 2.2 | <LOQ | 1.9 |
| 42 | 5.0 | 1.5 | <LOD | <LOD | 2.1 | <LOD | 1.8 |
| 43 | 5.1 | 1.2 | <LOD | <LOD | 2.7 | <LOQ | 2.3 |
| 44 | 4.7 | 1.0 | <LOD | <LOD | 2.2 | <LOD | 2.8 |
| 45 | 2.0 | 1.7 | <LOD | <LOD | 2.2 | <LOQ | 2.9 |
| 48 | 2.2 | 1.2 | <LOD | <LOD | 1.7 | <LOD | 2.2 |
| 49 | 1.4 | 1.0 | <LOD | <LOD | 1.6 | <LOD | 1.8 |
| 51 | 1.5 | 1.0 | <LOD | <LOD | 1.3 | <LOD | 1.8 |
| 54 | 2.3 | 1.1 | <LOD | <LOD | 1.5 | <LOD | 1.9 |
| 55 | 4.5 | 1.3 | <LOD | <LOD | 1.3 | <LOD | 2.2 |
| 56 | <LOD | 1.0 | <LOD | <LOD | 1.6 | <LOD | 1.8 |

**Table S5:** continued.

| | | | | | | | |
|---|---|---|---|---|---|---|---|
| 57 | 2.1 | 0.8 | <LOD | <LOD | 1.6 | <LOD | 2.1 |
| 59 | 1.9 | 1.2 | <LOD | <LOD | 1.9 | <LOD | 2.0 |
| 61 | 1.9 | 1.3 | <LOD | <LOD | 1.5 | <LOD | 2.1 |
| 65 | 1.8 | 1.2 | <LOD | <LOD | 1.5 | <LOD | 3.0 |
| 69 | 1.9 | 1.3 | <LOD | <LOD | 3.6 | <LOQ | 2.8 |
| 80 | 4.8 | 1.2 | <LOD | <LOD | <LOQ | <LOD | 2.9 |
| 87 | 4.1 | 1.2 | <LOD | <LOD | 3.4 | <LOQ | 2.1 |
| 89 | 1.6 | 1.1 | <LOD | <LOD | <LOQ | <LOD | 1.8 |
| 112 | 3.5 | 1.4 | <LOD | <LOD | 3.6 | <LOQ | 2.4 |
| 113 | 4.1 | 1.1 | <LOD | <LOD | 3.5 | <LOQ | 1.9 |
| 122 | 1.8 | 1.1 | <LOD | <LOD | 2.1 | <LOQ | 1.6 |
| 123 | 6.8 | 1.1 | <LOD | <LOD | 1.4 | <LOD | 1.9 |
| 126 | 2.2 | 1.0 | <LOD | <LOD | <LOQ | <LOD | 3.0 |
| 133 | 2.0 | 1.0 | <LOD | <LOD | 1.4 | <LOD | 1.6 |
| 136 | 3.8 | 1.3 | <LOD | <LOQ | 8.6 | 1.9 | <LOQ |
| 137 | 2.3 | 1.3 | <LOD | <LOD | 2.3 | <LOQ | 1.6 |
| 138 | 2.2 | 0.9 | <LOD | <LOD | 1.4 | <LOD | 1.7 |
| 140 | <LOD | 0.9 | <LOD | <LOD | 1.6 | <LOD | <LOQ |
| 141 | <LOD | 1.2 | <LOD | <LOD | 2.5 | <LOD | <LOQ |
| 142 | <LOD | 1.2 | <LOD | <LOD | 2.3 | <LOD | <LOQ |
| 145 | 2.9 | 1.1 | <LOD | <LOD | <LOQ | <LOD | <LOQ |
| 148 | 4.2 | 1.2 | <LOD | <LOD | 1.8 | <LOD | <LOQ |
| 152 | 3.9 | 1.2 | <LOD | <LOD | 1.7 | <LOD | <LOQ |
| 154 | 2.3 | 1.1 | <LOD | <LOD | 2.7 | <LOQ | 1.5 |
| 155 | <LOD | 1.5 | <LOD | <LOD | 1.7 | <LOD | 1.5 |
| 156 | 2.2 | 1.1 | <LOD | <LOD | 1.3 | <LOD | 2.5 |
| 157 | 1.6 | 1.4 | <LOD | <LOD | 2.6 | <LOQ | 1.5 |
| 159 | <LOD | 0.9 | <LOD | <LOD | 2.4 | <LOD | 2.1 |
| 160 | 1.9 | 1.0 | <LOD | <LOD | 2.3 | <LOQ | <LOQ |
| 161 | 2.5 | 1.3 | <LOD | <LOD | 2.4 | <LOQ | <LOQ |
| 162 | 2.7 | 1.3 | <LOD | <LOD | 2.8 | <LOQ | 2.1 |
| 164 | 2.1 | 1.5 | <LOD | <LOD | 2.7 | <LOQ | <LOQ |
| 165 | 12 | 1.0 | <LOD | <LOD | 1.9 | <LOQ | <LOQ |
| 166 | 2.7 | 1.3 | <LOD | <LOD | 2.4 | <LOQ | 1.8 |
| 167 | 33 | 1.1 | <LOD | <LOD | 1.7 | <LOQ | 1.5 |
| 168 | <LOD | 1.2 | <LOD | <LOD | 1.8 | <LOD | <LOQ |
| 170 | 2.0 | 1.6 | <LOD | <LOD | 1.3 | <LOD | <LOQ |
| 171 | 1.6 | 0.9 | <LOD | <LOD | 1.5 | <LOD | <LOQ |
| 173 | 2.0 | 1.2 | <LOD | <LOD | 1.4 | <LOD | <LOQ |
| 178 | 3.2 | 1.1 | <LOD | <LOD | 1.4 | <LOD | <LOQ |
| 181 | 2.0 | 1.0 | <LOD | <LOD | <LOQ | <LOD | <LOQ |
| 183 | <LOD | 1.7 | <LOD | <LOD | 3.0 | <LOQ | <LOQ |
| 211 | 1.6 | 1.4 | <LOD | <LOD | 1.7 | <LOQ | <LOQ |

[a] Limit of detection (LOD) and quantification (LOQ) according to Braun *et al.* (2020).
[b] Not assessed.
[c] Below the LOD.
[d] Below the LOQ.

**Table S6.** Concentration levels of mycotoxins found in breast milk samples obtained from three mothers on five consecutive days. All other 22 analytes were below the respective LOD value.

| **Analyte** | | **AME** | **BEA** | **EnnA** | **EnnA$_1$** | **EnnB** | **EnnB$_1$** | **OTA** |
|---|---|---|---|---|---|---|---|---|
| **Retention time (min)** | | 8.1 | 11.0 | 11.5 | 11.3 | 10.8 | 11.1 | 6.4 |
| **LOD (ng/L)[a]** | | 0.5 | 0.1 | 0.5 | 0.9 | 0.65 | 0.5 | 0.8 |
| **LOQ (ng/L)[a]** | | 1.0 | 0.3 | 1.0 | 1.8 | 1.3 | 1.0 | 1.5 |
| **Regression coefficient $R^2$** | | 0.999 | 0.999 | 0.999 | 0.999 | 0.999 | 0.999 | 0.999 |
| | | **AME** | **BEA** | **EnnA** | **EnnA$_1$** | **EnnB** | **EnnB$_1$** | **OTA** |
| **Volunteer** | **Day** | **(ng/L)** | | | | | | |
| **1** | 1 | 5.7 | 1.5 | <LOD[b] | <LOQ | 8.8 | 1.5 | 3.2 |
| | 2 | 3.6 | 2.2 | <LOD | <LOQ | 8.0 | 1.1 | 3.8 |
| | 3 | 8.1 | 1.6 | <LOD | <LOD | 4.9 | <LOQ[c] | 3.2 |
| | 4 | 3.9 | 1.2 | <LOD | <LOD | 4.7 | <LOQ | 2.0 |
| | 5 | 2.5 | 1.6 | <LOD | <LOD | 4.1 | <LOQ | 2.4 |
| **2** | 1 | 3.1 | 1.1 | <LOD | <LOD | 2.0 | <LOD | <LOD |
| | 2 | 11 | 0.9 | <LOD | <LOD | 2.0 | <LOD | <LOQ |
| | 3 | 1.8 | 0.9 | <LOD | <LOD | 1.7 | <LOD | <LOD |
| | 4 | <LOD | 1.0 | <LOD | <LOD | 2.3 | <LOD | <LOD |
| | 5 | <LOD | 0.8 | <LOD | <LOD | 1.8 | <LOD | 4.6 |
| **3** | 1 | 2.1 | 1.6 | <LOD | <LOD | 5.6 | <LOQ | 3.2 |
| | 2 | 12 | 1.1 | <LOD | <LOD | 2.8 | <LOQ | 3.5 |
| | 3 | 3.9 | 1.4 | <LOD | <LOD | 2.7 | <LOQ | 2.3 |
| | 4 | 1.6 | 1.5 | <LOD | <LOD | 4.7 | <LOQ | 3.0 |
| | 5 | 2.7 | 1.2 | <LOD | <LOD | 2.9 | <LOQ | 1.9 |

[a] Limit of detection (LOD) and quantification (LOQ) according to Braun *et al.* (2020).
[b] Below the LOD.
[c] Below the LOQ.

**Table S7.** Concentration levels of mycotoxins found in urine samples obtained from three mothers on five consecutive days. All other 20 analytes were below the respective limit of detection (LOD) value.

| **Analyte** | **AME** | **CIT** | **DON** | **OTA** | **ZEN** | **αZAL** | **αZEL** | **βZEL** |
|---|---|---|---|---|---|---|---|---|
| **Retention time (min)** | 20.7 | 19.0 | 8.5 | 20.6 | 20.8 | 19.6 | 19.8 | 19.0 |
| **LOD (ng/L)**[a] | 12 | 700 | 950 | 20 | 65 | 265 | 100 | 65 |
| **LOQ (ng/L)**[a] | 24 | 1400 | 1900 | 40 | 130 | 530 | 200 | 130 |
| **Regression coefficient $R^2$** | 0.999 | 0.998 | 0.999 | 0.998 | 0.999 | 0.999 | 0.999 | 0.999 |

| | | **AME** | **CIT** | **DON** | **OTA** | **ZEN** | **αZAL** | **αZEL** | **βZEL** |
|---|---|---|---|---|---|---|---|---|---|
| **Volunteer** | **Day** | **(ng/L)** | | | | | | | |
| | 1 | 132 | <LOD[b] | 45191 | 69 | 642 | 1848 | <LOQ[c] | <LOQ |
| | 2 | 73 | <LOQ | 87554 | 72 | 575 | 1124 | <LOQ | <LOQ |
| **1** | 3 | 49 | <LOD | 9017 | 51 | 242 | 599 | <LOQ | <LOD |
| | 4 | 33 | <LOQ | 11714 | <LOQ | 229 | 1130 | <LOQ | <LOD |
| | 5 | <LOD | <LOD | 12775 | 55 | <LOQ | <LOD | <LOD | <LOD |
| | 1 | 77 | <LOD | 8532 | <LOD | <LOQ | 2471 | <LOD | <LOD |
| | 2 | 37 | <LOD | 4087 | <LOD | <LOQ | 2331 | <LOD | <LOD |
| **2** | 3 | 100 | <LOD | 12859 | <LOQ | <LOQ | 2586 | <LOD | <LOD |
| | 4 | 72 | <LOD | 16493 | <LOD | 165 | 4756 | <LOD | <LOD |
| | 5 | 89 | <LOD | 21604 | 46 | 169 | 5228 | <LOD | <LOD |
| | 1 | 69 | <LOQ | 21270 | <LOQ | <LOD | <LOD | <LOD | <LOQ |
| | 2 | <LOD | <LOD | 11788 | 42 | <LOD | <LOD | <LOD | <LOD |
| **3** | 3 | <LOD | <LOD | 35301 | <LOD | <LOD | <LOD | <LOD | <LOD |
| | 4 | 44 | <LOQ | 10996 | 44 | <LOQ | <LOD | <LOD | <LOD |
| | 5 | 59 | <LOD | 9966 | <LOD | <LOD | <LOD | <LOD | <LOD |

[a] Limit of detection (LOD) and quantification (LOQ) as stated in Table S3.
[b] Below the LOD.
[c] Below the LOQ.
